# Right ventricular stiffening and anisotropy alterations in pulmonary hypertension: Mechanisms and relations to function

**DOI:** 10.1101/2024.05.24.592212

**Authors:** Sunder Neelakantan, Alexander Vang, Rana Raza Mehdi, Haley Phelan, Preston Nicely, Tasnim Imran, Peng Zhang, Gaurav Choudhary, Reza Avazmohammadi

## Abstract

**Aims:** Pulmonary hypertension (PH) results in an increase in RV afterload, leading to RV dysfunction and failure. The mechanisms underlying maladaptive RV remodeling are poorly understood. In this study, we investigated the multiscale and mechanistic nature of RV free wall (RVFW) biomechanical remodeling and its correlations with RV function adaptations.

**Methods and Results:** Mild and severe models of PH, consisting of hypoxia (Hx) model in Sprague-Dawley (SD) rats (n=6 each, Control and PH) and Sugen-hypoxia (SuHx) model in Fischer (CDF) rats (n=6 each, Control and PH), were used. Organ-level function and tissue-level stiffness and microstructure were quantified through in-vivo and ex-vivo measures, respectively. Multiscale analysis was used to determine the association between fiber-level remodeling, tissue-level stiffening, and organ-level dysfunction. Animal models with different PH severity provided a wide range of RVFW stiffening and anisotropy alterations in PH. Decreased RV-pulmonary artery (PA) coupling correlated strongly with stiffening but showed a weaker association with the loss of RVFW anisotropy. Machine learning classification identified the range of adaptive and maladaptive RVFW stiffening. Multiscale modeling revealed that increased collagen fiber tautness was a key remodeling mechanism that differentiated severe from mild stiffening. Myofiber orientation analysis indicated a shift away from the predominantly circumferential fibers observed in healthy RVFW specimens, leading to a significant loss of tissue anisotropy.

**Conclusion:** Multiscale biomechanical analysis indicated that although hypertrophy and fibrosis occur in both mild and severe PH, certain fiber-level remodeling events, including increased tautness in the newly deposited collagen fibers and significant reorientations of myofibers, contributed to excessive biomechanical maladaptation of the RVFW leading to severe RV-PA uncoupling. Collagen fiber remodeling and the loss of tissue anisotropy can provide an improved understanding of the transition from adaptive to maladaptive remodeling.

**Translational perspective:** Right ventricular (RV) failure is a leading cause of mortality in patients with pulmonary hypertension (PH). RV diastolic and systolic impairments are evident in PH patients. Stiffening of the RV wall tissue and changes in the wall anisotropy are expected to be major contributors to both impairments. Global assessments of the RV function remain inadequate in identifying patients with maladaptive RV wall remodeling primarily due to their confounded and weak representation of RV fiber and tissue remodeling events. This study provides novel insights into the underlying mechanisms of RV biomechanical remodeling and identifies the adaptive-to-maladaptive transition across the RV biomechanics-function spectrum. Our analysis dissecting the contribution of different RV wall remodeling events to RV dysfunction determines the most adverse fiber-level remodeling to RV dysfunction as new therapeutic targets to curtail RV maladaptation and, in turn, RV failure in PH.

## 1. Introduction

Pulmonary hypertension (PH) is defined as an elevation in mean pulmonary arterial pressure (mPAP) and is known to affect 1% of the world population ^1, 2^. The increase in RV afterload leads to increased RV wall stress, which causes alterations in the biomechanical behavior of the RV free wall (RVFW) myocardium ^3-6^ prominently defined by tissue stiffness and anisotropy. In patients with PH, disease severity and prognosis are drastically influenced by the ability of the RV to biomechanically adapt to increased afterload and maintain its function. Severe PH can lead to excessive changes to RVFW stiffness and anisotropy, and subsequently lead to RV failure ^7, 8^. However, the mechanisms behind changes in RVFW stiffness and anisotropy and their association with RV dysfunction remain inadequately studied.

The effect of passive remodeling on RV functional performance is also evidenced by a growing body of clinical and preclinical studies ^9^, including ours ^10-13^, suggesting that excessive stiffening of the RVFW correlates with worse outcomes in PH. However, it remains unclear as to when and how the stiffening becomes excessive and is considered maladaptive. RV fibrosis has been consistently observed in rodent models of PH ^14-16^, and it is commonly perceived as an increase in collagen content in the RVFW, leading to an increase in RVFW stiffness. However, recent studies of myocardial remodeling in both left and right heart diseases indicate that, in addition to the collagen content, changes in collagen fiber architecture are also an important contributor to overall biomechanical alterations of the myocardium and the chamber ^17-19^. In particular, collagen fiber undulation, which is inversely related to fiber tautness, is known to be a primary structural feature of collagen bundles in modulation of the passive stiffness of myocardial tissue. However, changes to the collagen fiber tautness, a primary characteristic of collagen architecture, and their effect on tissue stiffness in PH remain to be investigated. This is especially important as collagen fiber tautness can vary during collagen deposition (fibrosis), depending on diastolic stretching of the heart, leading to significantly different mechanical behavior at similar levels of fibrosis. The intricate interactions between cardiac remodeling at fiber, tissue, and organ levels necessitate a multiscale approach to understand the transition between adaptive to maladaptive RV in PH.

In addition to the level of RVFW stiffening, it is necessary to determine the directionality of the stiffening in response to PH. Directional stiffening combined with changes in the myofiber architecture can lead to changes in the anisotropy of the RVFW. The tissue anisotropy and underlying myofiber architecture of the myocardium ensure that the contractile behavior of the myofibers produces optimal active force and, subsequently optimal contractility to pump blood. Changes in tissue anisotropy can lead to significant changes in the directionality of the developed contractile forces that are predominantly circumferential in healthy RV myocardium. Indeed, changes in the directionality of the contractile forces alter the kinematics of contraction and can accelerate RV maladaptation and failure. Prior studies have reported changes in mean myofiber angle in the RVFW in PH ^19,20^. These studies indicated that adaptive architectural remodeling can improve RV function and that the myofiber angle could be an important prognostic tool during the assessment in PH. However, the change in myofiber angle also leads to changes in the anisotropy of the RV myocardium, affecting both the passive diastolic behavior and the active contractile behavior, including the force generation and wall motion. There is a lack of studies on the alteration in RVFW anisotropy and its effect on RV wall motion impairment, and consequently, RV function.

In this study, we present a multiscale investigation into the quantitative relationship between fiber-level remodeling and alteration in organ-level function using several distinct animal models of PH, providing a wide spectrum of RV remodeling, integrated with biomechanical modeling. We hypothesize that the reduction of RVFW tissue stiffening past a threshold leads to the transition from adaptive to maladaptive RV remodeling and, subsequently, RV dysfunction and RV failure. We performed mechanical tests to characterize the RVFW stiffening and changes in tissue anisotropy and studied the correlation between RVFW stiffness and hemodynamic indices. We performed multiscale biomechanical analysis to determine the level of contributions of fiber-level remodeling events such as myofiber reorientation, fibrosis, and increased collagen fiber tautness towards RVFW stiffness and anisotropy. Our findings provide novel insights into the mechanism of this stiffening and alterations in anisotropy, beyond the prevalent notion that RV stiffening is driven by an increase in RV mass or collagen content.

## 2. Methods

### 2.1 Characterizing RV dysfunction in PH through a spectrum of severity

All procedures were performed in accordance with the Guide for the Care and Use of Laboratory Animals published by the US National Institutes of Health. Approval for procedures on animal subjects was obtained from the Institutional Animal Care and Use Committee (IACUC) at the Providence VA Medical Center (IACUC 1633548). A mild model of PH was developed by placing Sprague Dawley (SD) rats (n=6) in a hypoxia chamber (10% O2) for three weeks (Hx). The corresponding wild-type control (CTL) rats (n=6) were placed in normoxia for the same duration. A severe model of PH was developed by injecting a vascular endothelial growth factor receptor inhibitor (VEGFR2 - SU5416) (20 mg/kg sc) into Fischer rats (CDF; n=6) and placing them in a hypoxia chamber for three weeks, followed by one week of normoxia (SuHx). The CTL rats (n=6) were placed in normoxia and were injected with vehicle instead of Sugen5416. All four groups had equal numbers of male and female rats used in the study ^21^. Additional SD control and SuHx rats (CTL, n=4; SuHx, n=4) were used to enhance the spectrum of RV adaptation to study the correlation between organ-level function and RVFW stiffening (reported in Fig. 2).

**Figure 1:**
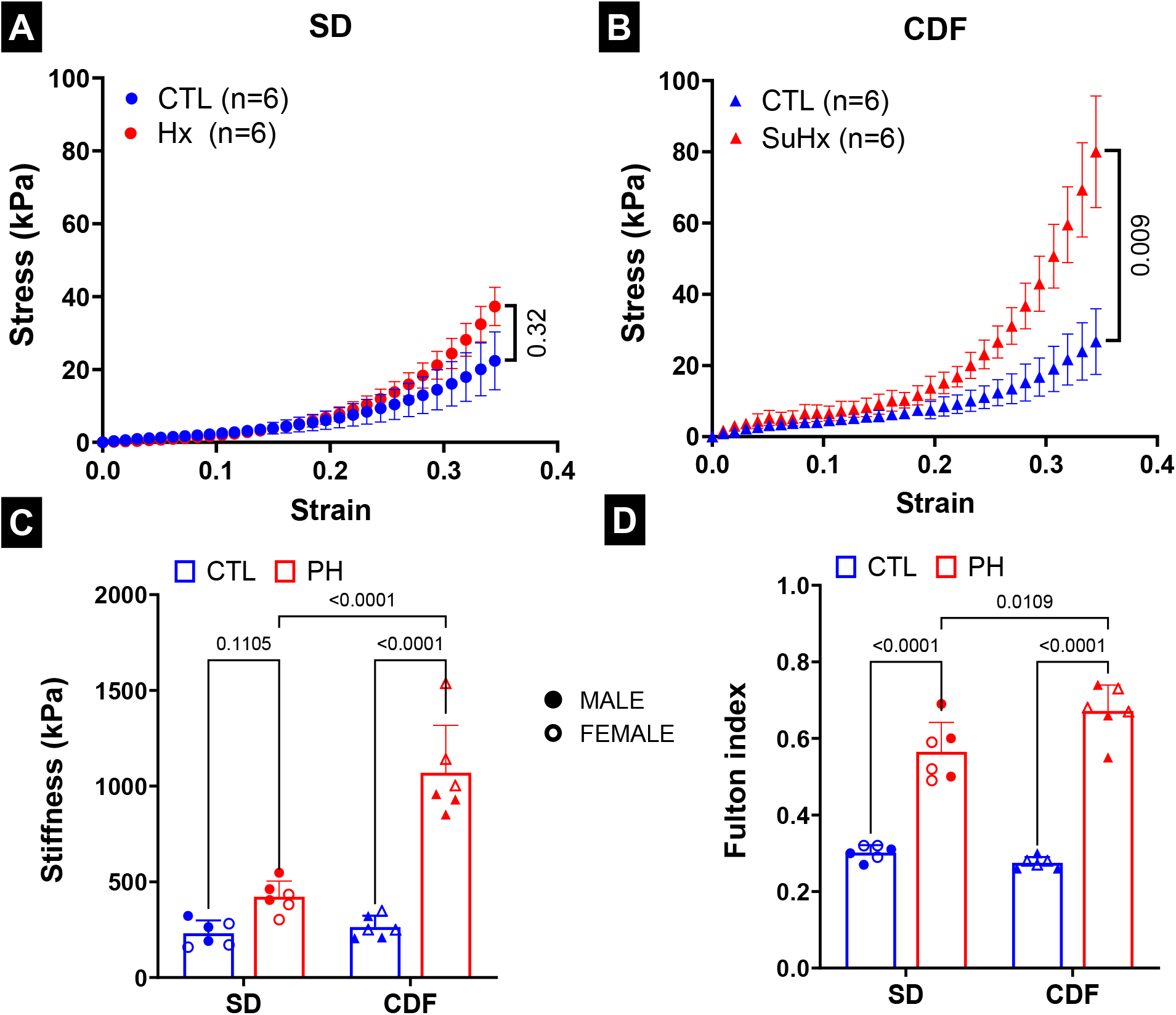
Right ventricle free wall (RVFW) stiffening and hypertrophy. Ensemble stress-strain plots obtain from biaxial testing of RVFW specimens from **(A)** SD and **(B)** CDF rats. Ensemble stress = Circumferential + Longitudinal stress. **(C)** Stiffness obtained as the slope of the stress-strain plots at 35% strain. **(D)** Fulton Index. The filled and hollow markers indicate male and female specimens, respectively. Statistical significance in **(A)** and **(B)** was calculated using unpaired student t-tests. Statistical significance in **(C)** and **(D)** was calculated using 2-way ANOVA with Tukey’s multiple comparisons. SD: n=6 CTL, 6 PH (Hx); CDF: n=6 CTL, 6 PH (SuHx).

**Figure 2:**
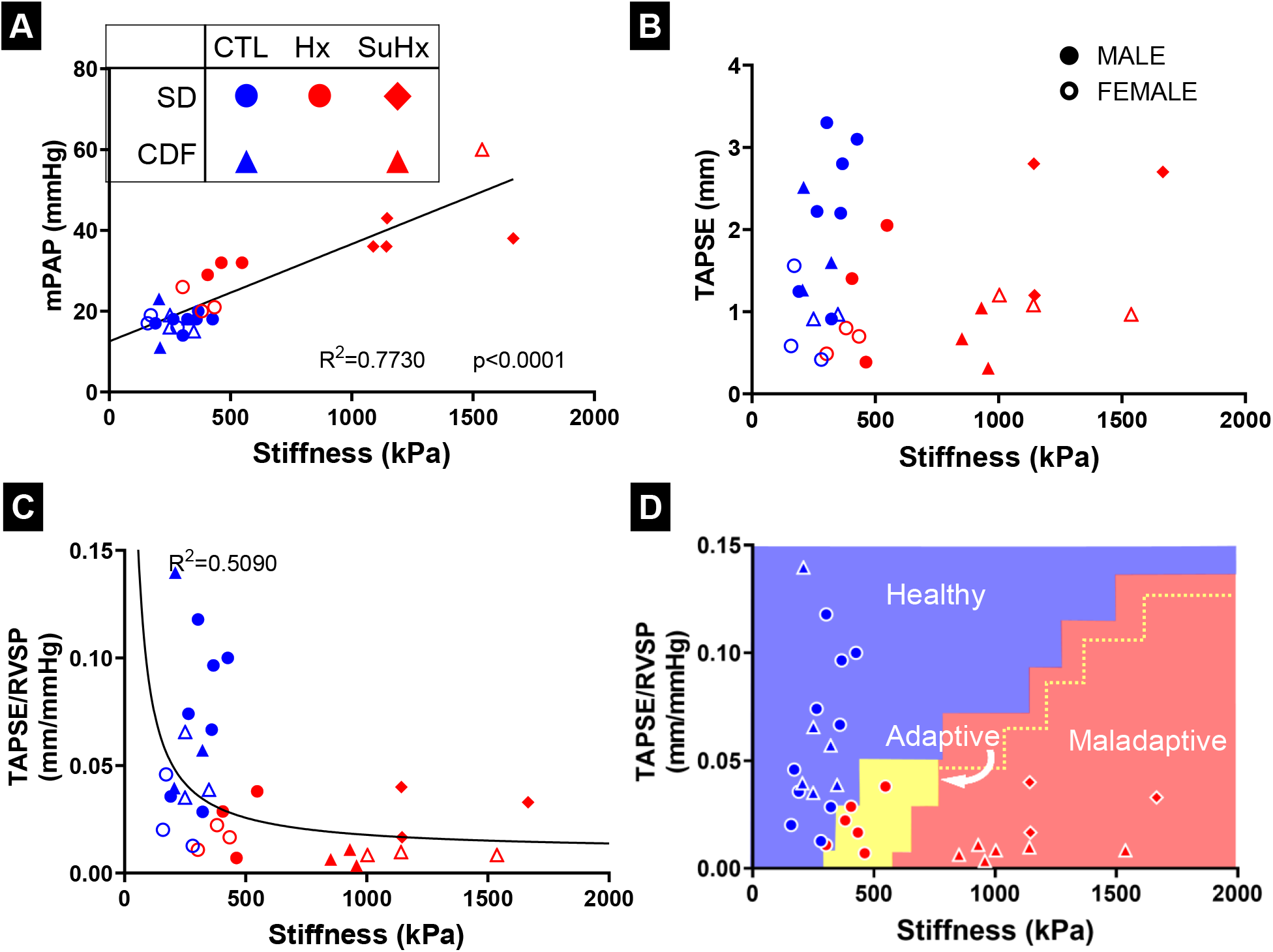
Association between tissue-level stiffening and organ-level dysfunction. Correlation between stiffness and **(A)** mean pulmonary arterial pressure (mPAP), **(B)** TAPSE and **(C)** TAPSE/RVSP. **(D)** Thresholding between adaptive and maladaptive regions estimated through k-nearest neighbor classification. Yellow dotted line indicates the expected extension of the adaptive region. The filled and hollow markers indicate male and female specimens, respectively. Linear regression performed in **(A)**, and significance is reported for non-zero slope. SD: n=6 CTL, 6 PH (Hx); SD: n=4 CTL, 4 PH (SuHx); CDF: n=6 CTL, 6 PH (SuHx).

### 2.2 Characterizing RV dysfunction in PH through in vivo measurements

At the end of the study, animals were placed on a heating pad (37°C) and anesthetized with continuous isoflurane inhalation (1.5%–2%) in 100% O_2_ for the duration of the echocardiography and catheterization procedures. Transthoracic echocardiography was performed as previously described ^14^. Briefly, a linear array rodent transducer was used on the Vevo 2100 (FUJIFILM VisualSonics, Toronto, ON, Canada) to obtain echocardiographic measurements. Pulmonary acceleration time (PAT) was measured using the pulsed-wave Doppler recording at the RV overflow tract. Tricuspid annular plane systolic excursion (TAPSE) was determined by measuring the excursion of the tricuspid annulus from its highest position to the peak descent during ventricular systole, using M-mode echocardiography. The early diastolic velocity of the RV lateral wall (at tricuspid annulus) was measured using Tissue Doppler echocardiography. The Tissue Doppler was also used to measure the tricuspid valve (TV) early wave peak (TVE) and TV atrial wave peak (TVA).

Post echocardiography, catheterization was performed using an open-chest technique and a high-fidelity Millar 2.0F catheter (SPR-869). The catheter was inserted into the RV through the apex to measure RV systolic pressure (RVSP) and end-diastolic pressure (RVEDP). The catheter was then placed into the pulmonary artery to measure mPAP. The RV-PA coupling was determined through the parameter TAPSE/RVSP.

### 2.3 Biaxial testing to measure the tissue-level RV stiffening and anisotropy

After catheterization, the animals were euthanized under isoflurane anesthesia by excising the heart for tissue collection. After the animals were sacrificed, the hearts were excised from the chest cavity. The atria were removed, and RVFW was isolated from the heart. The weights of the RVFW and remaining ventricular myocardium were recorded to determine the Fulton index, defined as the ratio of RVFW weight to the combined weight of the LVFW and septum.

After isolation of the RVFW, the tissue was trimmed into a rectangular slab with the sides being aligned along the anterior-posterior, apex-base, and transmural directions. The RVFW thickness (RVFWT) was measured for each specimen using calipers. The slabs were mounted in a biaxial mechanical testing machine (Cellscale Biotester, CellScale, Waterloo, Ontario, Canada) along circumferential and longitudinal directions, respectively (Fig. S1-I). The myocardium specimens were submerged in the phosphate-buffered saline at 22°C during testing to prevent dehydration. Testing consisted of 10 cycles of preconditioning to equalize all residual stresses in the tissue, followed by 10 cycles of equibiaxial stretch, where each direction was stretched by 30%. The stress in this study was calculated as the ratio of force over an initial cross-sectional area. The stiffness along each direction was calculated as the slope of the stress-strain curve at 30% strain (Fig. S1-I). This stiffness was summed together to report an effective stiffness for the corresponding RVFW tissue, which will be referred to as RVFW stiffness.

In addition to tissue stiffness, tissue anisotropy was calculated as the ratio of the circumferential to longitudinal mechanical response. Given this definition, two metrics of the anisotropy were presented: (i) stiffness anisotropy, calculated as the ratio of circumferential to longitudinal stiffness (i.e., the ratio of the slopes to the respective stress-strain curves), and (ii) stress anisotropy, calculated as the ratio of circumferential to longitudinal stress (i.e., the ratio of the respective stress values). Both metrics were calculated at 35% strain. These anisotropy metrics are expected to reflect the effective myo- and collagen fiber architecture in the RVFW. After characterizing the organ-level function and tissue-level mechanical response, we investigated the correlation between tissue-level biomechanical remolding, such as changes in stiffness and anisotropy, and hemodynamic adaptations using regression analysis.

### 2.4 Histological analysis to determine the fiber-level alteration in RV architecture

After mechanical testing, all the RVFW specimens were fixed in 10% formalin for 48 hours and stored in 70% ethanol. The specimens were sectioned transmurally with a slice thickness of 5 µm every 200 µm and stained with picrosirius red (PSR) stain, turning myofibers yellow and collagen fibers red (Fig. S1-II). These specimens were mounted on slides and then imaged using the Olympus VS120 Virtual Slide Scanning System to obtain images at 20x magnification. The images were used to calculate the distribution of myofiber angles as a function of transmural depth using methods described in our previous work ^22-24^. Briefly, the Beta distribution functions of myofiber angles were calculated at each depth level and stacked together to obtain the fiber distribution as a function of transmural depth. In addition to myofiber and collagen fiber orientation, the images were used to calculate the collagen content. Stack of images for each RVFW specimen were processed to separate myofiber pixels (P_*c*_) and collagen fiber pixels (P_*m*_), and the myofiber and collagen fiber contents were estimated as Φ_*m,c*_ = P_*m,c*_/(P_*c*_ + P_*m*_).

The histological slides were also used to quantify the collagen content to determine the degree of fibrosis. We characterized collagen fiber tautness through a tautness metric, defined as the ratio of the distance between the endpoints of collagen fibers and their arc lengths (Fig. S1-II). A value closer to 1 indicates a straight/taut fiber, and a value close to 0 indicates a fully coiled fiber.

### 2.5 Separating contributions of fiber-level remodeling events to stiffening

We used a constitutive model, developed to describe the biomechanical behavior of the myocardium ^25-27^, to separate the contribution of myo- and collagen fiber towards the tissue-level stiffness. Briefly, the overall energy function was defined as

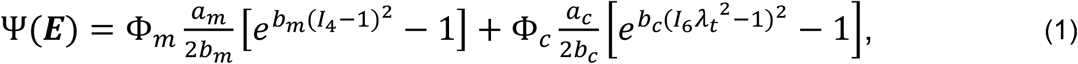

where*a*_*m*_, *b*_*m*_ are the material constants characterizing the myofiber behavior; *a*_*c*_, *b*_*c*_, *λ*_*s*_ are the material constants characterizing the collagen behavior, with the collagen fiber tautness being characterized by *λ*_*t*_ (*λ*_*t*_ = 1/*λ*_*s*_, where *λ*_*s*_ denotes the stretch required to remove the slack in the collagen fibers; Φ_*m*_ and Φ_*c*_ represent the myo- and collagen-fiber volume fractions in the tissue specimen; and *I*_4_ *I*_6_ are invariants characterizing the deformation of the myofiber and collagen fibers, respectively. For the optimization process, myofiber and collagen content, transmural fiber orientation, and collagen fiber undulation were measured from histology (detailed in section 2. 4) and incorporated into Eq. 1. Also, the exponential terms in the myofiber and collagen components (*b*_*m*_, *b*_*c*_) were fixed to ensure uniqueness in the optimization. Thus, all changes in stiffness were manifested in *a*_*m*_ and *a*_*c*_ that were obtained using Marquette-Levenberg least-square optimization.

### 2.6 Statistical analysis

The data in this study was analyzed in GraphPad Prism 9 and Microsoft Excel. The data were represented as mean ± standard deviation. Both male and female rats were included in the study and have been represented using solid and hollow markers, respectively. For correlation plots, goodness of fit (R^2^) and the significance of non-zero slope were used to characterize the linear fits. Non-linear regression analyses were characterized by goodness of fit (R^2^). K-nearest neighbor classification (scikit-learn, python) was used to identify adaptive and maladaptive RVFW stiffening regions. Unpaired student t-tests were used when comparing the control and PH samples of the same strain/ severity. When comparing all four groups, we used two-way ANOVA with Tukey’s correction. Significance was accepted at p<0.05. We have chosen to report the numerical values of significance.

## 3. Results

### 3.1 PH caused organ-level dysfunction and tissue-level hypertrophy and stiffening

A significant increase was observed in RVSP in both Hx (mild) and SuHx (severe) models of PH (Table 1). In contrast, RVEDP increased significantly only in SuHx rats, but not in Hx rats. TAPSE decreased significantly in the SuHx rats, leading to a significant decrease in RV-PA coupling (TAPSE/RVSP) (Table 1). The trend was similar in Hx rats, but the change was not significant (Table 1).

**Table 1.** Hemodynamic measurements of both severities of PH, separated by strain and PH severity. Data represented as mean ± standard deviation. SD: n=6 CTL, 6 PH (Hx); CDF: n=6 CTL, 6 PH (SuHx). Significance was reported using unpaired students’ t-tests. BPM – beats per minute, RVSP – RV systolic pressure, RVEDP – RV end-diastolic pressure, TAPSE – tricuspid annular plane systolic excursion, PAT – Pulmonary acceleration time, TV E – tricuspid valve (TV) E wave velocity, TVA – TV A-wave velocity, RVFWT – RVFW thickness.

| Catherization |  |  |  |  |
| --- | --- | --- | --- | --- |
|  | Strain | Control | PH | Significance |
| Heart rate (BPM) | SD | 371.48±61.59 | 372.66±29.25 | 0.967 |
| | CDF | 365.30±39.69 | 351.27±20.67 | 0.496 |
| RVSP (mmHg) | SD | 32.17±2.32 | 46.83±7.31 | <0.001 |
| | CDF | 26.33±4.76 | 110.48±18.70 | <0.001 |
| RVEDP (mm Hg) | SD | 5.93±0.83 | 6.78±1.20 | 0.185 |
| | CDF | 6.33±0.82 | 10.53±4.14 | 0.035 |
| TAPSE/RVSP (mm/mmHg) | SD | 0.036± 0.022 | 0.021± 0.012 | 0.154 |
| | CDF | 0.063± 0.040 | 0.008± 0.003 | 0.007 |
| Echocardiography |  |  |  |  |
|  | Strain | Control | PH | Significance |
| Cardiac output (ml/min) | SD | 63.56±14.36 | 50.08±15.46 | 0.149 |
| | CDF | 36.90±15.25 | 14.30±6.11 | 0.013 |
| LV ejection fraction (%) | SD | 83.70±8.87 | 86.62±7.35 | 0.548 |
| | CDF | 89.26±7.88 | 97.68±5.05 | 0.070 |
| PAT (s) | SD | 32.07±2.63 | 28.67±7.84 | 0.337 |
| | CDF | 24.77±6.31 | 13.86±1.38 | 0.002 |
| TAPSE (mm) | SD | 1.16±0.67 | 0.97±0.64 | 0.634 |
| | CDF | 1.53±0.61 | 0.88±0.33 | 0.046 |
| TV A (mm/s) | SD | 656.36±217.35 | 669.61±274.70 | 0.931 |
| | CDF | 545.08±98.80 | 341.60±357.22 | 0.24 |
| TV E (mm/s) | SD | 407.77±107.28 | 400.57±163.19 | 0.932 |
| | CDF | 248.31±62.87 | 536.52±142.55 | 0.01 |
| RVFWT (mm) | SD | 0.45±0.08 | 0.81±0.25 | 0.006 |
| | CDF | 0.46±0.19 | 1.20±0.26 | <0.001 |

Exponential-like stress-strain behavior of the RVFW showed upward stiffening in both Hx and SuHx rats for combined strain, but the change was significant only for SuHx rats (Figs. 1A,B). Both longitudinal and circumferential strains were significantly increased in the SuHx rats (Figs. S2A,B). These changes were similarly reflected in the RVFW stiffness measured at strain=0.35 (Fig. 1C). In addition, the Fulton index increased significantly in both strains (Fig. 1D), and the increase agreed with the RVFW thickening measured from echocardiography (Table 1). While significant RVFW hypertrophy was noted in both Hx and SuHx rats, RVFW stiffening was present only in SuHx rats. This suggested additional RVFW stiffening mechanisms beyond RVFW thickening, as detailed in sections 3.4 and 3.5.

### 3.2 Tissue-level RVFW stiffening was associated with RV afterload and organ-level RV dysfunction

RVFW stiffness showed strong positive correlations with mPAP (Fig. 2A, R^2^ =0.77, p <0.01) and RVSP (Fig. S3A, R^2^ =0.68, p <0.01). RVFW stiffness also correlated with RVEDP and PAT (Figs. S3B,C). However, no observable correlation between RVFW stiffness and TAPSE could be detected (Fig. 2B). In contrast, RV-PA coupling (measured as TAPSE/RVSP) and RVFW stiffness demonstrated an inverse hyperbolic-like relationship, indicating that accelerating increases in RVFW stiffness could lead to significant decreases in RV-PA coupling (Fig. 2C). K-nearest neighbor classification corroborated this observation, resulting in smaller and larger adaptive and maladaptive regions, respectively, with a step-like interface between all the regions (Fig. 2D). The dashed yellow line indicates the expected extension of the adaptive region in case additional data covering a wider range of RV-PA coupling and RVFW stiffening is present.

### 3.3 PH led to the loss of myofiber anisotropy in the RV myocardium

Healthy RVFW specimens exhibited a strong circumferential bias (significantly higher stiffness in the circumferential direction), while RVFW specimens from the CDF SuHx rats did not (Fig. 3A). This decrease in circumferential bias was also observed in the anisotropy ratio of the SuHx rats (Fig. S4). TAPSE/RVSP showed significant correlations with changes in RVFW anisotropy (Fig. 3B).

**Figure 3:**
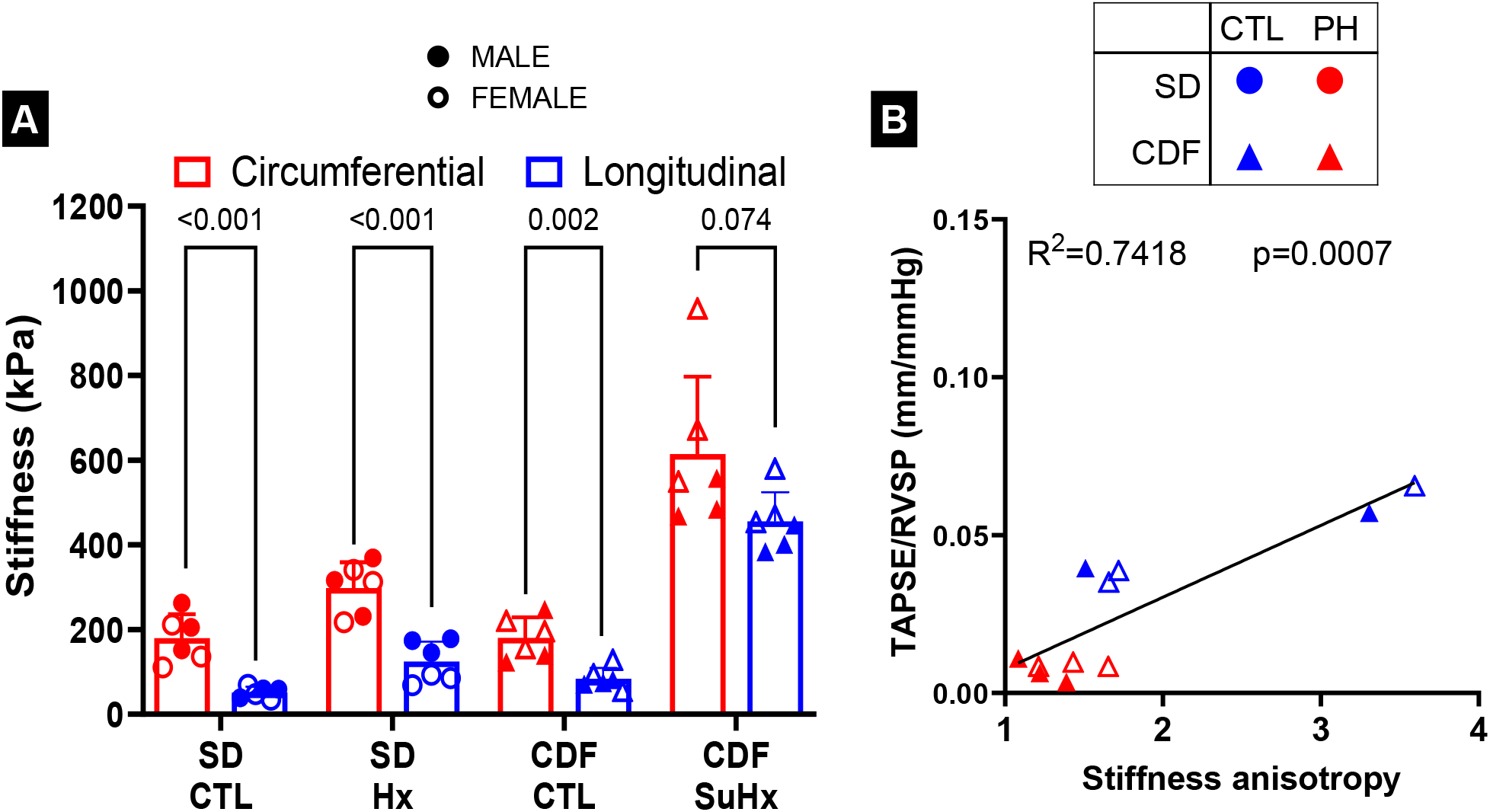
RVFW Anisotropy and association with RV function. **(A)** Anisotropy of the RVFW specimens estimated by comparing the statistical significance between circumferential and longitudinal stiffness. **(B)** Correlation between anisotropy and TAPSE/RVSP for the CDF specimens. The filled and hollow markers indicate male and female specimens, respectively. Statistical significance in **(A)** was calculated using unpaired student t-tests. Linear regression performed in **(B)** and significance is reported for non-zero slope. SD: n=6 CTL, 6 PH (Hx); CDF: n=6 CTL, 6 PH (SuHx).

### 3.4 PH-induced fiber-level changes to myocardial architecture

There was a shift in the myofiber angle away from the circumferential direction in the mid-section of the RVFW in both groups of PH, with the change being significant only for SuHx rats (Fig. S5). This observation confirmed the changes in tissue anisotropy seen in the ex-vivo biaxial testing (Figs. 4, S5). This change in myofiber angle also manifested in the slope of the myofiber angle as a function of transmural thickness. The absolute value of the slope decreased in Hx rats and increased in SuHx rats (Figs. 4A,B), indicating that fibers tend to orient towards circumferential direction in mild PH (Hx rats) but longitudinal direction in severe PH (SuHx rats). In contrast, the myofiber helical range did not change in either group (Fig. 4C). The changes in myofiber orientation were concentrated near the endocardial region in Hx rats, whereas the changes were more uniform in SuHx rats (Fig. S5).

**Figure 4:**
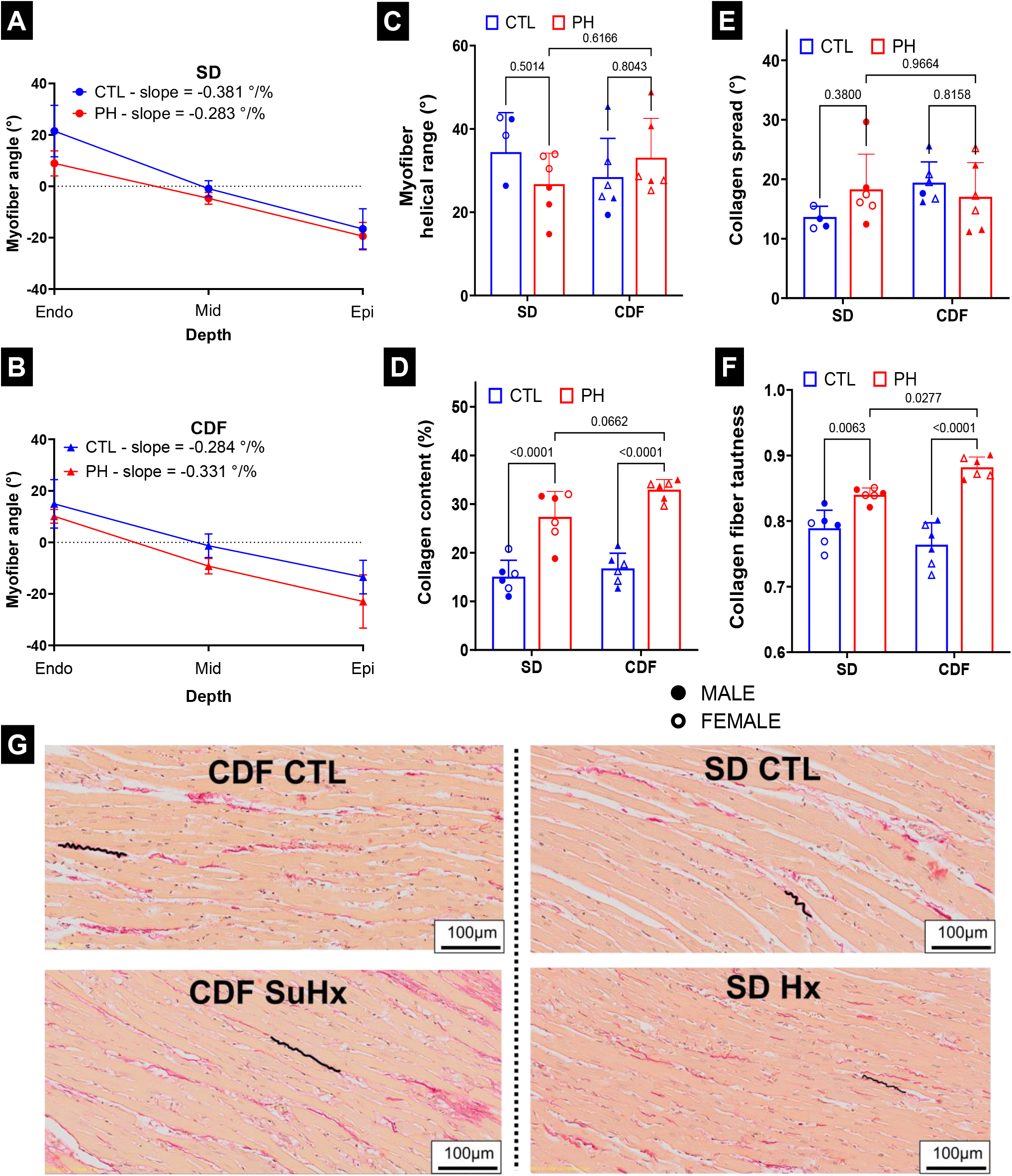
Architectural remodeling of right ventricle. Variation of myofiber angle with transmural depth in **(A)** SD and **(B)** CDF rats. **(C)** Myofiber helical range. **(D)** Collagen fiber content, **(E)** collagen fiber spread and **(F)** Collagen fiber tautness (tautness=1 indicates fully straight/taut fibers). **(G)** Histological images of the RV specimens marked with representative collagen fibers to indicate tautness. The filled and hollow markers indicate male and female specimens, respectively. Statistical significance in **(C), (D), (E)** and **(F)** was calculated using 2-way ANOVA with Tukey’s multiple comparison. SD: n=5-6 CTL, 6 PH (Hx); CDF: n=6 CTL, 6 PH (SuHx).

In contrast to the mild changes in the myofiber architecture, the collagen content increased in both severities of PH, indicating significant fibrosis (Fig. 4D) in both models. The collagen fiber spread (the standard deviation of collagen fiber angle within a transmural layer) remained similar in healthy and PH rats (Fig. 4E), suggesting that the increased collagen deposition took place along the direction of myofibers (Fig. S1(II)). Importantly, although collagen fiber tautness increased in both severities of PH, the increases in SuHx rats were significantly more than that in Hx rats (Figs. 4F,G).

### 3.5 Collagen fiber tautness significantly contributed to tissue-level stiffness

Through constitutive modeling, we estimated the contribution of myofiber and collagen fibers toward the tissue-level stiffness. The overall myofiber stiffness (combining volume fraction and intrinsic fiber stiffness) and *intrinsic* myofiber stiffness remained unchanged (Fig. 5A and 5B). In contrast, the overall collagen fiber stiffness increased significantly in SuHx rats (Fig. 5C), while fiber-level stiffness remained unchanged (Fig. 5D). The multiscale observations comprising (i) RVFW stiffened in SuHx group, but not Hx (Fig. 1D), (ii) collagen content similarly increased in both groups (Fig. 3D), and (iii) intrinsic collagen fiber stiffness remained unchanged in both groups (Fig. 5D), suggested that collagen content is insufficient to determine the level of stiffening and that the effect of collagen fiber tautness should also be incorporated when dissecting the mechanisms contributing to RVFW stiffening. The constitutive model (Eq. 1) indeed precisely deconvolutes the effect of collagen content (Φ_*c*_), collagen fiber stiffness (*a*_*c*_, *b*_*c*_), and collagen fiber tautness (*λ*_*t*_) on RVFW tissue-level behavior. Overall collagen stiffness (a_*c*_Φ_*c*_) correlated with the RVFW stiffness of each severity (Hx and SuHx) (Fig. 6A). Notably, the SuHx rats had a steeper relationship in comparison to the Hx rats, which is consistent with the higher tautness in SuHx rats (Fig 4F). Indeed, collagen fiber tautness (reduced fiber slack) showed a steep exponential-like association with increased RVFW stiffness (Fig. 6B), whic was predicted by the constitutive model (Eq. 1).

**Figure 5:**
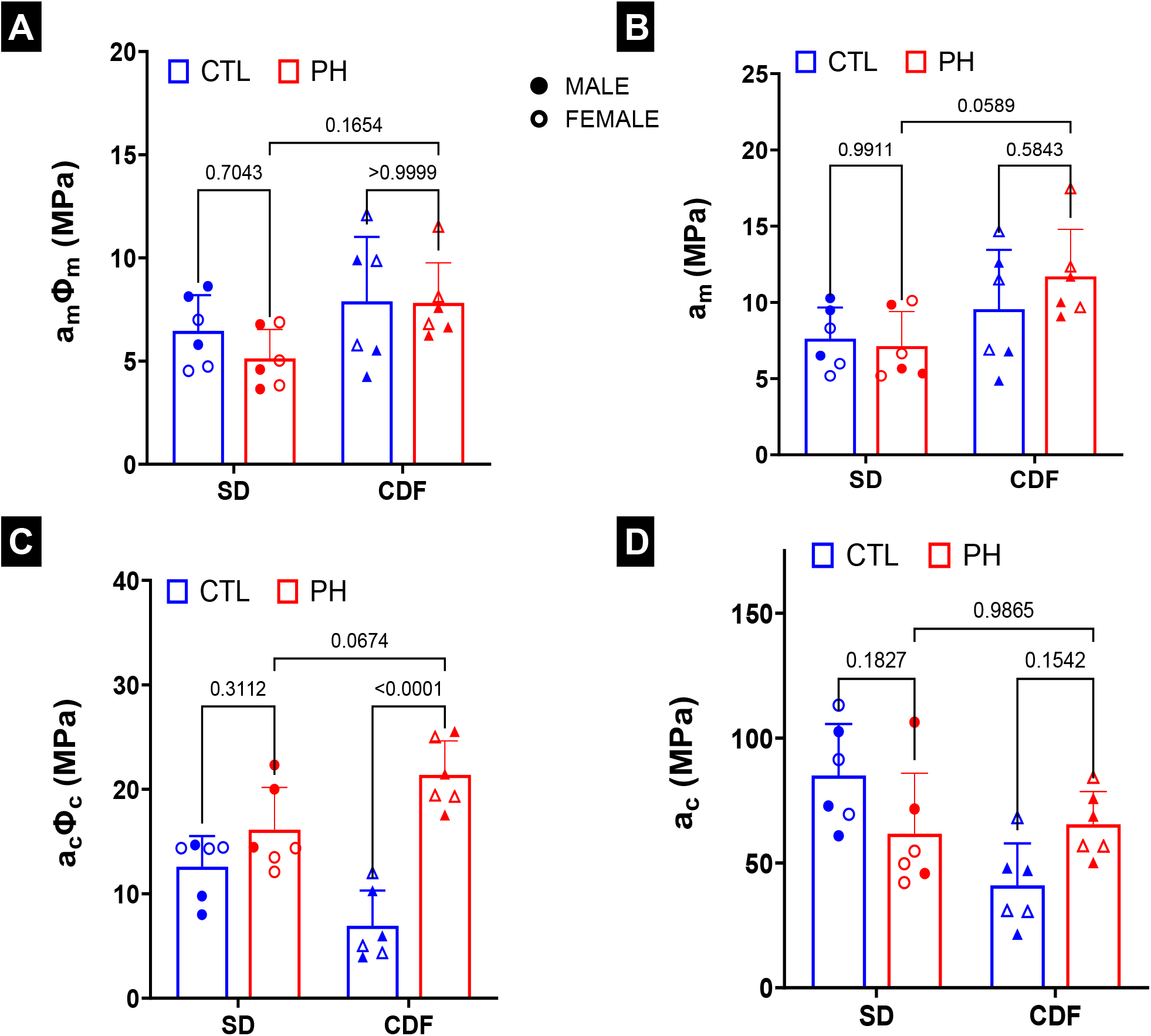
Contributions from fiber-level remodeling towards RVFW stiffening. Stiffness parameters were obtained from fitting stress-strain data to a material model of the myocardium. **(A)** overall and **(B)** fiber-level myofiber stiffness. **(C)** overall and **(D)** fiber-level collagen fiber stiffness. The filled and hollow markers indicate male and female specimens, respectively. Statistical significance in **(A), (B), (C)** and **(D)** was calculated using 2-way ANOVA with Tukey’s multiple comparison. SD: n=6 CTL, 6 PH (Hx); CDF: n=6 CTL, 6 PH (SuHx).

**Figure 6:**
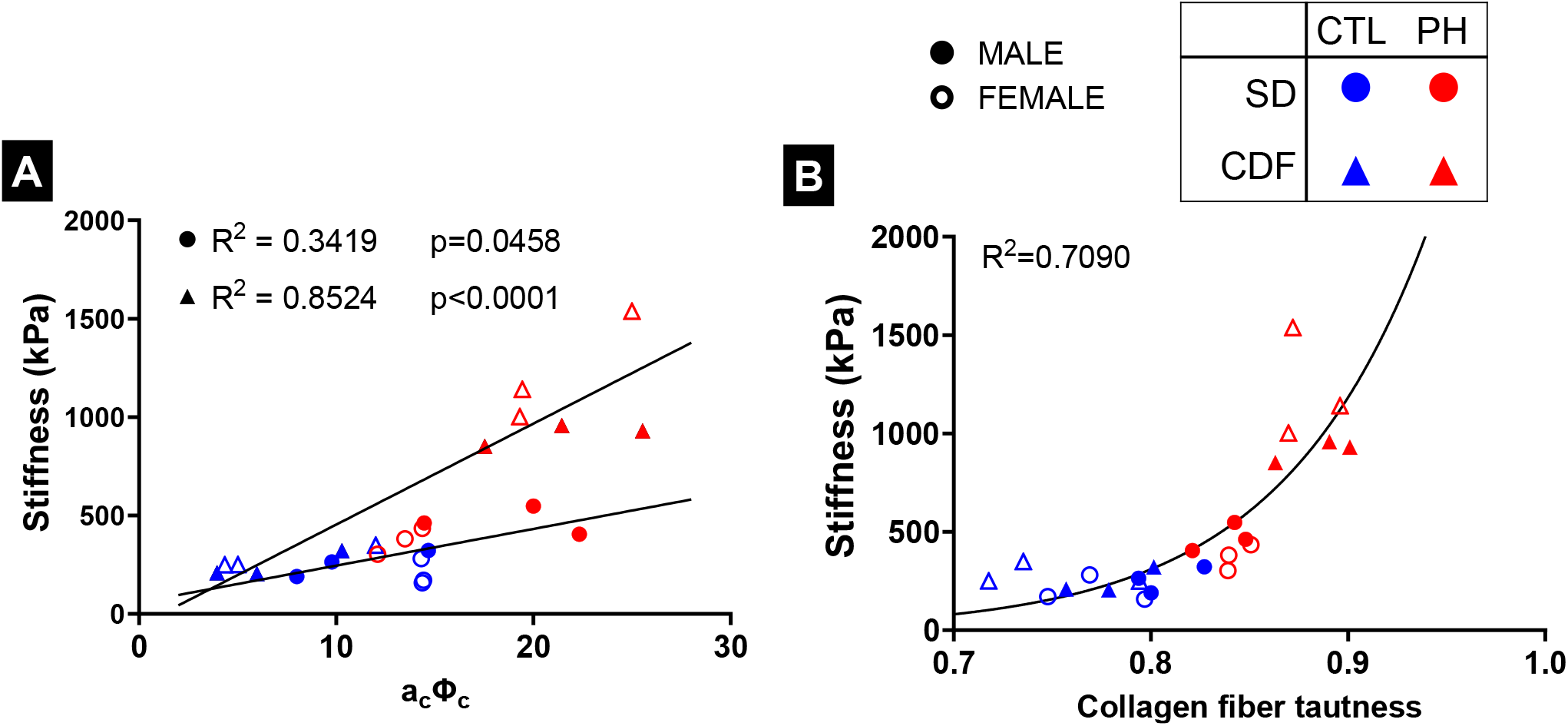
Association between collagen fiber remodeling and RVFW stiffening. **(A)** Association between RVFW stiffness normalized by collagen fiber content and collagen fiber stiffness. **(B)** Association between RVFW stiffness and collagen fiber tautness. The filled and hollow markers indicate male and female specimens, respectively. SD: n=6 CTL, 6 PH (Hx); CDF: n=6 CTL, 6 PH (SuHx).

## 4. Discussion

### 4.1 Mild and severe PH models enabled the dissection of RVFW remodeling events driving RV maladaptation

In this study, we investigated the stiffening and loss of anisotropy of the RVFW myocardium in PH, their effect on RV dysfunction, and their underlying mechanisms using Hx and SuHx rodent models in PH. While RVSP and RVFWT increased significantly in both Hx and SuHx rats, several hemodynamic parameters such as RVEDP, TAPSE, PAT, and TAPSE/RVSP increased significantly only in SuHx rats. This allowed us to establish a spectrum of PH severity from healthy to mild PH to severe PH. Through this spectrum, we investigated the variation of RVFW stiffness and tissue anisotropy with PH severity. We quantified the contribution of key mechanisms underlying tissue-level stiffening and loss of anisotropy, including hypertrophy, myofiber reorientation, collagen fibrosis, and increase in collagen fiber tautness, and their association with RV-PA uncoupling as a function of PH severity. These analyses enabled us to predict the adaptive and maladaptive domains of RV remodeling in terms of the severity of RVFW biomechanical remodeling.

### 4.2 RVFW stiffening level differentiated adaptive and maladaptive RV remodeling

The RVFW stiffness increased in both severities of PH, with the increase being significant only in SuHx rats. Stiffness was strongly correlated with RVSP and PAT, and inversely correlated with RV-PA coupling (TAPSE/RVSP). The strong correlation of RVFW stiffness with RVSP and mPAP suggests that the RVFW stiffening is primarily driven by elevations in RV afterload in systole. In addition, RVFW stiffening promotes an increase in RVEDP as increased stiffness reduces myocardial passive stretching in diastole. Additionally, PAT decreased in PH, and the decrease was linearly correlated with RVFW stiffening. Reductions in PAT are likely caused by the decrease in pulmonary artery (PA) compliance, as expected from a fluid mechanics perspective^28^. The decrease in PA compliance is corroborated by the correlation recently observed between RVFW stiffening and stiffening in PA in the mild (Hx) and severe (SuHx) rodent models of PH^29^. Moreover, the hyperbolic association between RVFW stiffness and RV-PA coupling indicated a strong non-linear nature of the adaptive-to-maladaptive transition of RV dysfunction. Using the k-nearest neighbor classification method, we found a narrow “buffer” range of RVFW stiffening where RV-PA coupling is maintained and beyond which the RV accelerates to maladaptation manifested by significant RV-PA uncoupling.

### 4.3 Severe PH led to a reduction in RVFW anisotropy

In addition to increased stiffness, the anisotropy of the RVFW specimens changed, with the biaxial test results indicating RVFW tends to transition towards “isotropy” in PH. The change in tissue anisotropy can significantly change the contractile behavior of the RVFW by causing a shift in the direction of the contractile forces produced by the myofibers. This change will directly affect the pumping efficiency of the RV at the organ level, even if the magnitude of contractile forces remains the same at the fiber level. Healthy RVFW specimens were strongly anisotropic, with a stiffer response in the circumferential direction, supported by the myofiber orientation distribution. However, the RVFW specimens in PH rats tended to reorient toward the longitudinal direction, leading to a reduced contrast between the circumferential and longitudinal directions. This phenomenon can be interpreted as a more isotropic behavior, evidenced by the anisotropy metric moving towards one. Alterations in the RVFW anisotropy that were estimated from biaxial testing were in agreement with changes in the myofiber orientation, indicating that fibers move away from the circumferential direction, which was pronounced in the mid-section of the myocardium in SuHx rats. Indeed, alterations in myofiber orientation serve as the primary mechanism of changes in anisotropy reduction.

The transition of RVFW biomechanics towards an isotropic behavior has also been reported in other decompensated rodent models of PH, including PA banding (PAB) and monocrotaline models ^6,30,31^, and is consistent with “sphericalization” of the RV at very high systolic pressure. Interestingly, the hyperbolic correlation between RVSP and anisotropy suggests that the transition towards an isotropic RVFW begins as a compensatory mechanism at moderate elevations of RVSP, providing a fair balance in active stress elevations between longitudinal and circumferential directions. In contrast, extreme reduction in anisotropy at high RVSP in SuHx rats leads to RV decompensation due to significant suppression in circumferential contractility ^19^, which is the prevalent mode of contractility for the healthy RV. As a result, the anisotropy metric showed a significant correlation with RV-PA coupling in SuHx rats, although the correlation was close to significant once combined with Hx rats.

### 4.4 Changes in collagen fiber architecture was a key contributor to RVFW stiffening

Our multiscale stress-strain modeling differentiated between the increase in RVFW stiffness occurring due to increased fiber content versus changes in intrinsic architectural and material properties of individual fibers. The intrinsic myofiber stiffness increased mildly in SuHx rats (insignificant) and the intrinsic myofiber stiffness in Hx rats remained unchanged. Studies have noted increases in isolated RV cardiomyocytes passive tension in human patients with pulmonary arterial hypertension ^32^ and in rodent PH model by PAB^33^. However, unchanged passive tension in isolated RV cardiomyocytes in human patients with class-II PH (caused by left heart failure) was also reported ^34^. We performed comparisons with the results by Rain et al.^33^, who studied the contribution of myofibrils and collagen towards passive RVFW behavior through ex-vivo testing. This study reported mild increases in myofiber contribution to RV passive stress in a severe PAB model of PH in male Wistar rats (Fig. 1B in ^33^). To perform proper comparisons, we zeroed the contribution of collagen fibers in Eq. 1 and created stress-strain plots for myofiber contribution only (Figs. S6A, B). The estimated myofiber tension was mildly increased in the SuHx model (Fig. S6B), which agrees with findings in ^33^ (Fig. 1B in ^33^).

In contrast to myofiber stiffness, the overall collagen fiber stiffness increased significantly in both severities of PH. However, similar to the myofiber, the individual collagen fiber stiffness did not increase significantly, indicating that a relative increase in collagen content versus myofiber (of nearly the same stiffnesses) contributed to the RVFW stiffening. On the other hand, similar increases in the collagen content were observed for both severities, begging the question of the contribution of additional remodeling events to excessive stiffening in SuHx rats. Interestingly, the RVFW stiffness (normalized by collagen content) in SuHx rats showed a much stronger association with the fiber-level collagen stiffness in comparison to that in Hx rats. This observation indicated that similar increases in collagen content contributed very differently to RVFW stiffness in Hx and SuHx rats, resulting in a significantly stiffer RVFW in the SuHx rats. This difference was linked to the collagen fiber architecture being significantly tauter in SuHx rats, which was also corroborated by a steep exponential-like association between RVFW stiffness and collage tautness. This alteration in the collagen fiber architecture has been a rather overlooked mechanism of RV stiffening in PH. Alterations in collagen fiber tautness have also been observed in other structural heart diseases ^10,18,35^ and in diseases associated with other soft connective tissues containing collagen ^36-39^. These studies indicate that changes in collagen fiber tautness could notably contribute to alterations in tissue stiffness and are *independent* of changes in collagen content. This mechanism explained the RVFW stiffening not only at high strains but also at low strains where collagen contribution to tissue stiffness is known to be minimal compared to myofibers due to their undulated nature (Fig. S7). Physiologically, the RVFW is under low strains during diastole, where the myocardium is expected to easily stretch at low pressures. RVFW stiffening at low strains (due to de-slacking of collagen fibers; Fig. S7) leads to restrictive filling during diastole. Therefore, the RV requires an increase in diastolic pressure to maintain proper filling in PH. This proposed mechanism is supported by the observed RVEDP values from the severe PH model (Table 1).

Overall, when investigating the mechanisms contributing to alterations in RVFW stiffness and anisotropy, the key contributors that differentiated adaptive from maladaptive remodeling were myofiber reorientation and increases in collagen fiber tautness. In contrast, low-fidelity makers such as Fulton index remained insensitive and confounded in clearly distinguishing adaptive from maladaptive remodeling. Our findings call for a mechanistic approach in characterizing the severity of RV remodeling in PH that accounts for *intrinsic* RVFW stiffness and fiber-level contribution to the stiffness. Quantification of these higher-fidelity markers of RV remodeling remains a possible route that could be sought by integrating advanced medical imaging, hemodynamic measurements, and digital twin creation.

### 4.5 Limitations

The RVFW passive stiffening, changes in the anisotropy, and architectural remodeling of the RVFW also affect the contractile behavior of the RV. We have presented some preliminary results in a recent study ^40^, and we will perform a comprehensive characterization of RVFW contractile adaptations and their relationship with RVFW stiffening and RV dysfunction in future studies. Previous studies reported that sex is an important factor in the incidence of PH ^41,42^, with the female-to-male ratio of incidence at 4.3:1 for pulmonary arterial hypertension ^43^. Future studies will investigate the effect of sex on the tissue-level RV remodeling behavior observed in this study, such as fibrosis and stiffening, and their subsequent effect on RV function in various severities of PH. Last, validation of the RV remodeling mechanisms identified in our preclinical study will be pursued in control and PH human cohorts in future studies.

## 5. Conclusion

In this study, we investigated the relationship between RVFW stiffening and loss of anisotropy with (mal)adaptation of RV function in PH. Our study indicates that tissue-level myocardial stiffening and loss of anisotropy play an important role in the alteration of RV function. Our multiscale biomechanical analysis, integrated with in-vivo and ex-vivo studies of mild and severe models of PH, identified key fiber remodeling mechanisms driving changes in RVFW biomechanics and, in turn, RV decompensation. The assimilation of multi-rodent model data led to quantitatively identifying the adaptive and maladaptive regions across the RV biomechanics-function domain which remains a key knowledge gap in understanding the onset and progression of RV remodeling in PH. The new mechanistic insights into the adaptive-to-maladaptive RV transition presented in this study, integrated with advances in RVFW imaging, promise to provide a significant departure from inadequate global assessments of RV remodeling, leading to improved diagnosis, prognosis, and therapeutics in PH patients.

## 7. Acknowledgements

We would like to acknowledge the assistance of the Integrated Microscopy and Imaging Laboratory at the Texas A&M College of Medicine. Some of the material of this work was supported with resources and the use of facilities at the VA Providence Healthcare System and the CardioPulmonary Vascular Biology COBRE (under National Institute of General Medical Sciences grant P20GM103652).

## Supplementary Material

**Figure S1:**
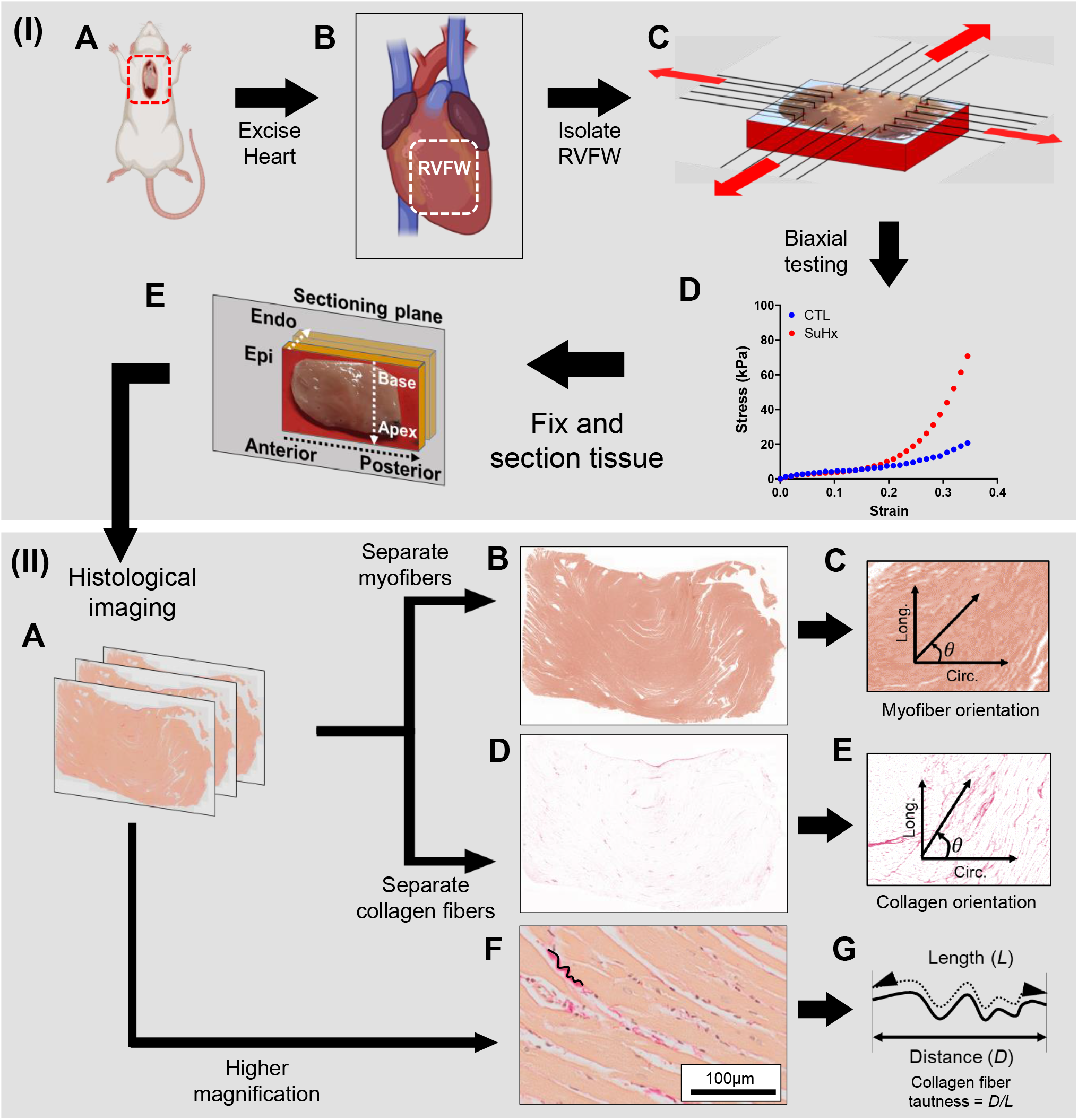
Schematic of the ex-vivo methods used in this study. (**I**) **(A)** Excision of heart from rat and **(B)** isolation of right ventricle free wall tissue (RVFW) specimen, followed by **(C**,**D)** ex-vivo biaxial testing and **(E)** fixation and sectioning of tissue. (**II**) Histological analysis of fixed tissue. Tissues were **(A)** imaged at two magnifications. The lower magnification images were separated into **(B)** myo- and **(D)** collagen fibers and used to **(C**,**E)** calculate the respective fiber orientation. **(F)** The higher magnification images were used to estimate collagen fiber slack using the equation in **(G)**.

**Figure S2:**
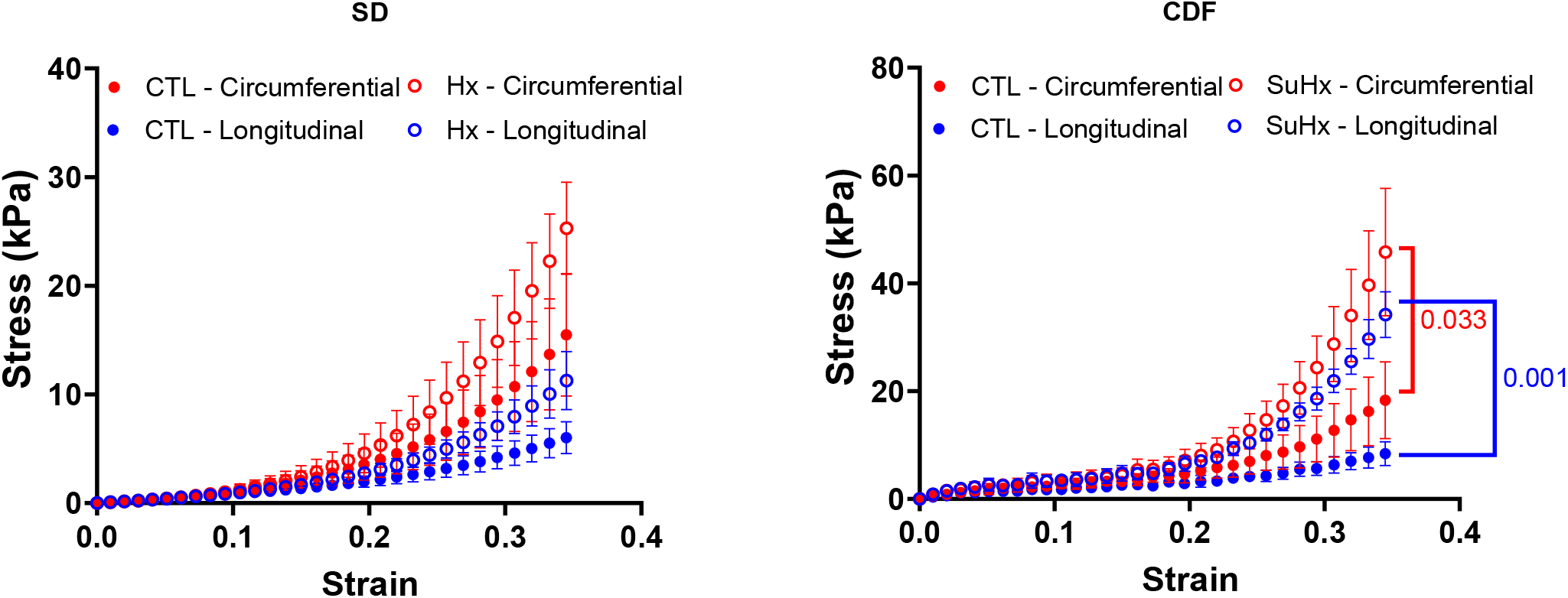
Biaxial testing results separated by strain and sex. **(A)** CDF rats and **(B)** SD rats. Statistical significance was calculated using unpaired student t-tests SD: n=6 WT, 6 PH (Hx); CDF: n=6 CTL, 6 PH (SuHx).

**Figure S3:**
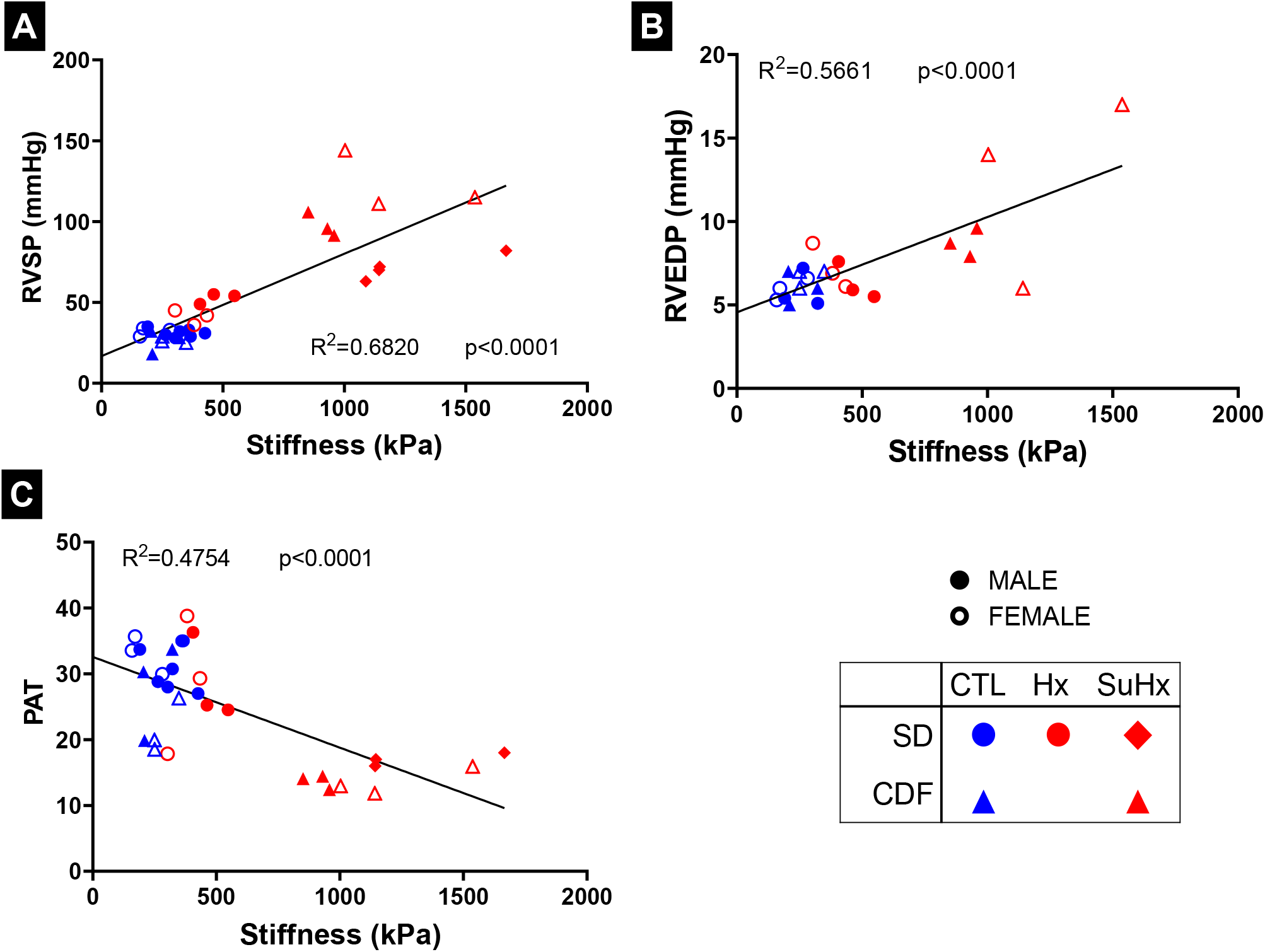
Correlation between RVFW stiffness and **(A)** RVSP, **(B)** RVEDP, and **(C)** pulmonary artery acceleration time (PAT). The filled and hollow markers indicate male and female specimens, respectively. Linear regression was performed in **(A)**-**(C)** significance is reported for non-zero slope. SD: n=10 WT, 6 PH (Hx), 4 PH (SuHx); CDF: n=6 CTL, 6 PH (SuHx).

**Figure S4:**
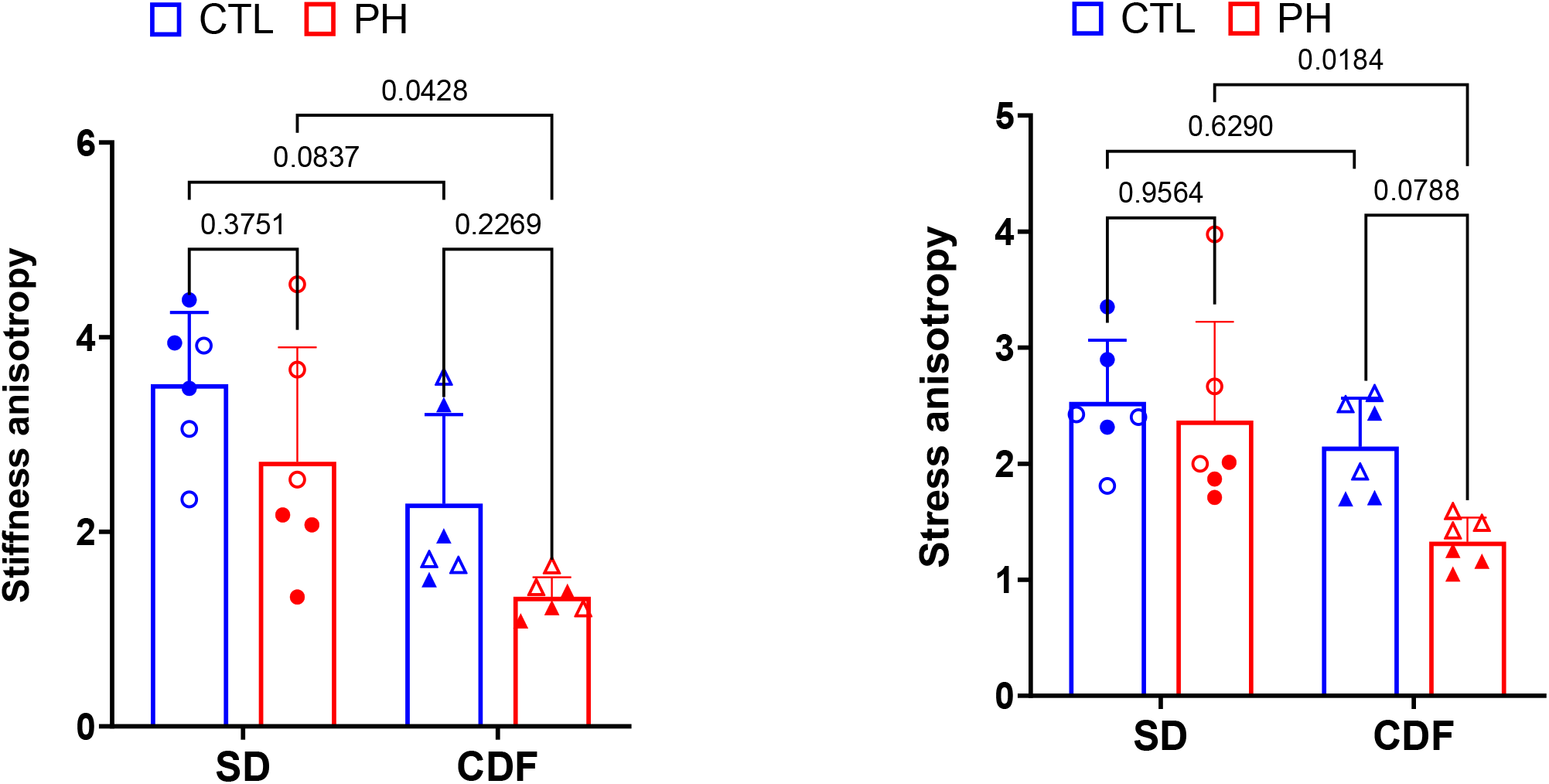
Anisotropy, calculated as the ratio of circumferential to longitudinal **(A)** stiffness and **(B)** maximum stress. The filled and hollow markers indicate male and female specimens, respectively. Statistical significance was calculated using 2-way ANOVA with Tukey’s multiple comparison. SD: n=6 CTL, 6 PH (Hx); CDF: n=6 CTL, 6 PH (SuHx).

**Figure S5:**
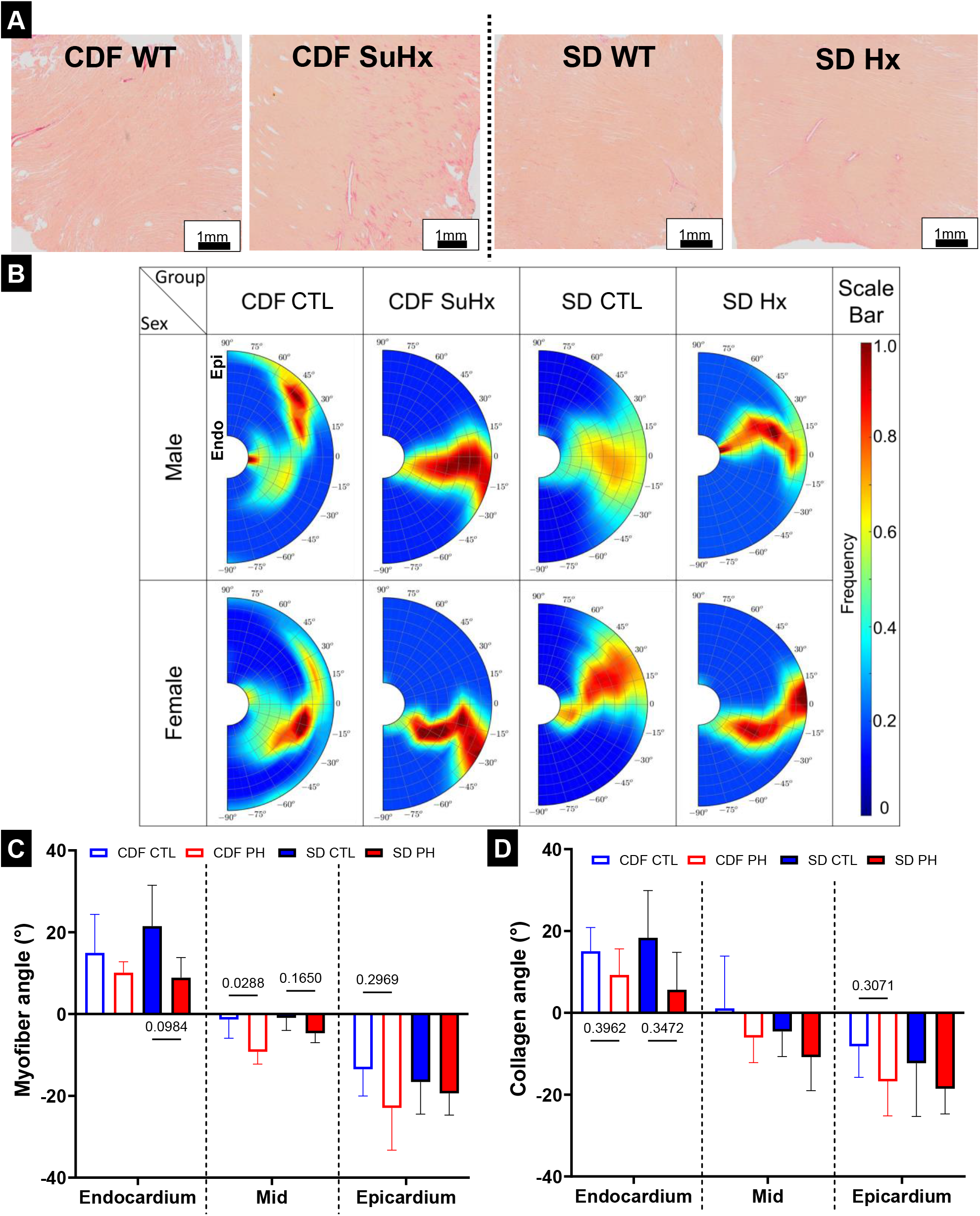
**(A)** Representative histological images of each group of rats used in the study. **(B)** Representative transmural myofiber orientation distribution. **(C)** Myofiber orientation at the endocardium, mid myocardium and epicardium. **(D)** Collagen fiber orientation at the endocardium, mid myocardium and epicardium.. Statistical significance in **(C)** and **(D)** was calculated using 2-way ANOVA with Tukey’s multiple comparison. SD: n=6 CTL, 6 PH (Hx); CDF: n=6 CTL, 6 PH (SuHx).

**Figure S6:**
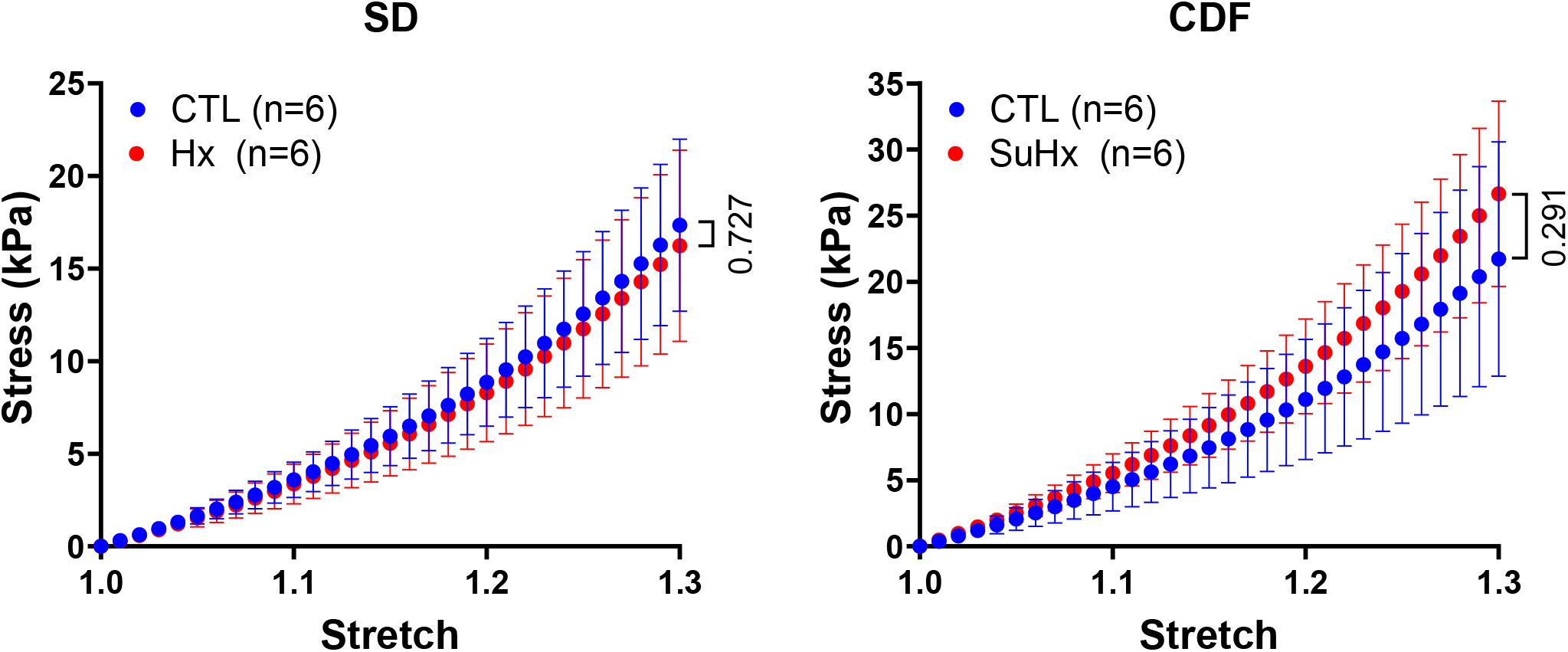
Estimated stress-stretch behavior of only the myofiber component of the **(A)** SD (CTL and Hx) rats, and **(B)** CDF (CTL and SuHx) rats. Stretch is defined as the ratio of current to the original dimension of the tissue. Statistical significance were calculated using unpaired student t-tests. SD: n=6 CTL, 6 Hx; CDF: n=6 CTL, 6 SuHx.

**Figure S7:**
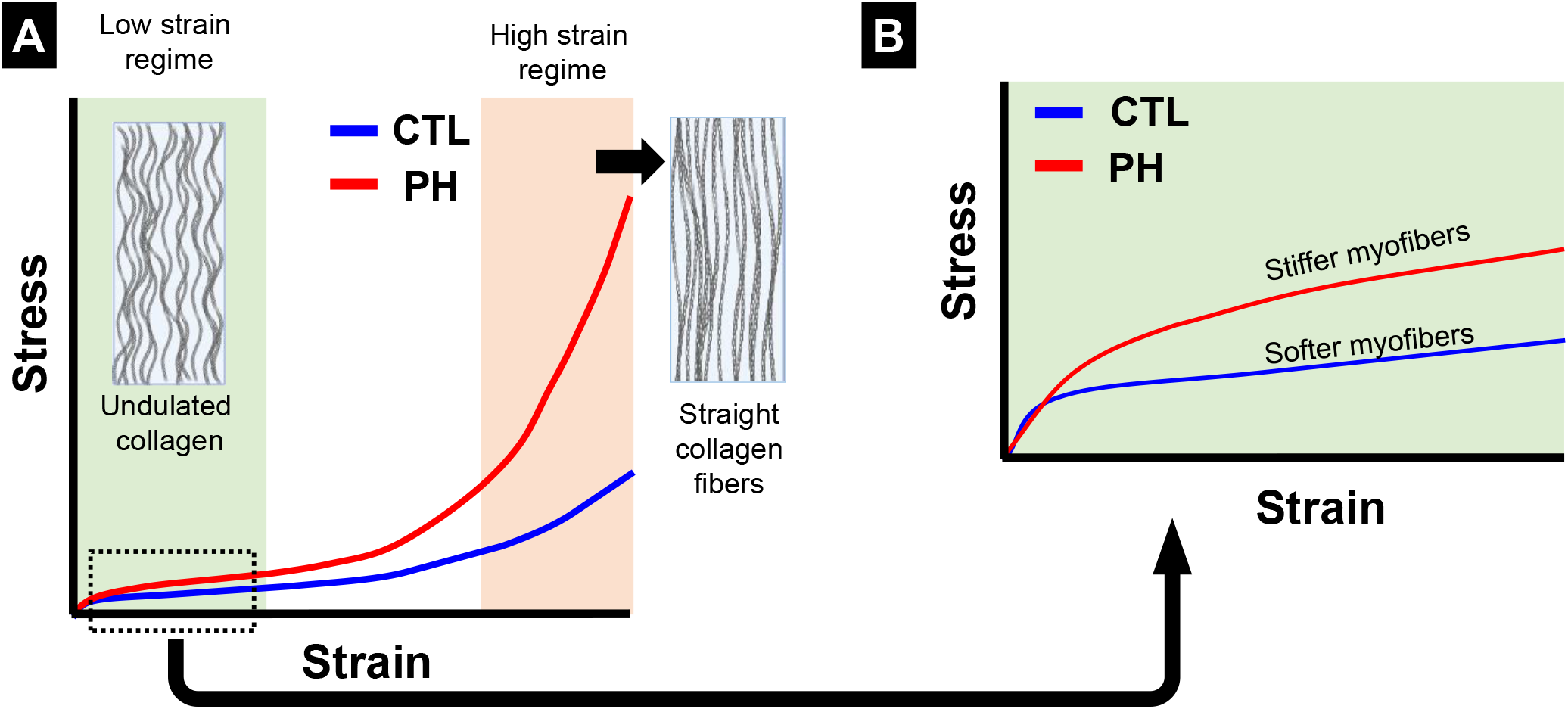
**(A)** Schematic of stress-strain behavior of myocardial tissue **(B)** Stress-strain behavior focused on low strains where the contribution from collagen is expected to be minimal..

## References

1. Hoeper, M. M.; Ghofrani, H. A.; Grunig, E.; Klose, H.; Olschewski, H.; Rosenkranz, S., Pulmonary Hypertension. Dtsch Arztebl Int 2017, 114 (5), 73–83.

2. Naeije, R.; Manes, A., The right ventricle in pulmonary arterial hypertension. Eur Respir Rev 2014, 23 (134), 476–487.

3. Bernardo, R. J.; Haddad, F.; Couture, E. J.; Hansmann, G.; Perez, V. A. D.; Denault, A. Y.; de Man, F. S.; Amsallem, M., Mechanics of right ventricular dysfunction in pulmonary arterial hypertension and heart failure with preserved ejection fraction. Cardiovasc Diagn The 2020, 10 (5), 1580–1603.

4. Vonk-Noordegraaf, A.; Haddad, F.; Chin, K. M.; Forfia, P. R.; Kawut, S. M.; Lumens, J.; Naeije, R.; Newman, J.; Oudiz, R. J.; Provencher, S.; Torbicki, A.; Voelkel, N. F.; Hassoun, P. M., Right heart adaptation to pulmonary arterial hypertension: physiology and pathobiology. Turk Kardiyol Dern A 2014, 42, 29–44.

5. Franco, V., Right Ventricular Remodeling in Pulmonary Hypertension. Heart Fail Clin 2012, 8 (3), 403-+.

6. Hill, M. R.; Simon, M. A.; Valdez-Jasso, D.; Zhang, W.; Champion, H. C.; Sacks, M. S., Structural and Mechanical Adaptations of Right Ventricle Free Wall Myocardium to Pressure Overload. Annals of Biomedical Engineering 2014, 42 (12), 2451–2465.

7. Ryan, J. J.; Archer, S. L., The Right Ventricle in Pulmonary Arterial Hypertension Disorders of Metabolism, Angiogenesis and Adrenergic Signaling in Right Ventricular Failure. Circ Res 2014, 115 (1), 176–188.

8. Voelkel, N. F.; Quaife, R. A.; Leinwand, L. A.; Barst, R. J.; McGoon, M. D.; Meldrum, D. R.; Dupuis, J.; Long, C. S.; Rubin, L. J.; Smart, F. W.; Suzuki, Y. J.; Gladwin, M.; Denholm, E. M.; Gail, D. B., Right ventricular function and failure - Report of a National Heart, lung, and Blood Institute working group on cellular and molecular mechanisms of right heart failure. Circulation 2006, 114 (17), 1883–1891.

9. Hayabuchi, Y.; Ono, A.; Homma, Y.; Kagami, S., Analysis of Right Ventricular Myocardial Stiffness and Relaxation Components in Children and Adolescents With Pulmonary Arterial Hypertension. J Am Heart Assoc 2018, 7 (9).

10. Mendiola, E. A.; Neelakantan, S.; Xiang, Q.; Merchant, S.; Li, K.; Hsu, E. W.; Dixon, R. A. F.; Vanderslice, P.; Avazmohammadi, R., Contractile Adaptation of the Left Ventricle Post-myocardial Infarction: Predictions by Rodent-Specific Computational Modeling. Annals of Biomedical Engineering 2023, 51 (4), 846–863.

11. Avazmohammadi, R.; Mendiola, E. A.; Soares, J. S.; Li, D. S.; Chen, Z. Q.; Merchant, S.; Hsu, E. W.; Vanderslice, P.; Dixon, R. A. F.; Sacks, M. S., A Computational Cardiac Model for the Adaptation to Pulmonary Arterial Hypertension in the Rat. Annals of Biomedical Engineering 2019, 47 (1), 138–153.

12. Avazmohammadi, R.; Mendiola, E. A.; Li, D. S.; Vanderslice, P.; Dixon, R. A. F.; Sacks, M. S., Interactions Between Structural Remodeling and Hypertrophy in the Right Ventricle in Response to Pulmonary Arterial Hypertension. J Biomech Eng-T Asme 2019, 141 (9).

13. Avazmohammadi, R.; Hill, M.; Simon, M.; Sacks, M., Transmural remodeling of right ventricular myocardium in response to pulmonary arterial hypertension. Apl Bioengineering 2017, 1 (1).

14. Vang, A.; Bos, D. D. G.; Fernandez-Nicolas, A.; Zhang, P.; Morrison, A. R.; Mancini, T. J.; Clements, R. T.; Polina, I.; Cypress, M. W.; Jhun, B. S.; Hawrot, E.; Mende, U.; O-Uchi, J.; Choudhary, G., alpha 7 Nicotinic acetylcholine receptor mediates right ventricular fibrosis and diastolic dysfunction in pulmonary hypertension. Jci Insight 2021, 6 (12).

15. Simpson, C. E.; Hassoun, P. M., Myocardial Fibrosis as a Potential Maladaptive Feature of Right Ventricle Remodeling in Pulmonary Hypertension. Am J Resp Crit Care 2019, 200 (6), 662–663.

16. Andersen, S.; Nielsen-Kudsk, J. E.; Noordegraaf, A. V.; Man, F. S. d., Right Ventricular Fibrosis. Circulation 2019, 139 (2), 269–285.

17. Emilio, A. M.; Sunder, N.; Qian, X.; Shuda, X.; Jianyi, Z.; Vahid, S.; Peter, V.; Reza, A., An image-driven micromechanical approach to characterize multiscale remodeling in infarcted myocardium. Acta Biomaterialia 2023.

18. Torres, W. M.; Barlow, S. C.; Moore, A.; Freeburg, L. A.; Hoenes, A.; Doviak, H.; Zile, M. R.; Shazly, T.; Spinale, F. G., Changes in Myocardial Microstructure and Mechanics With Progressive Left Ventricular Pressure Overload. Jacc-Basic Transl Sc 2020, 5 (5), 463–480.

19. Mendiola, E. A.; Bos, D. D. G.; Leichter, D. M.; Vang, A.; Zhang, P.; Leary, O. P.; Gilbert, R. J.; Avazmohammadi, R.; Choudhary, G., Right Ventricular Architectural Remodeling and Functional Adaptation in Pulmonary Hypertension. Circ-Heart Fail 2023, 16 (2), 202–211.

20. Kia, D. S.; Benza, E.; Bachman, T. N.; Tushak, C.; Kim, K.; Simon, M. A., Angiotensin Receptor-Neprilysin Inhibition Attenuates Right Ventricular Remodeling in Pulmonary Hypertension. J Am Heart Assoc 2020, 9 (13).

21. Cheron, C.; McBride, S. A.; Antigny, F.; Girerd, B.; Chouchana, M.; Chaumais, M.-C.; Jaïs, X.; Bertoletti, L.; Sitbon, O.; Weatherald, J.; Humbert, M.; Montani, D., Sex and gender in pulmonary arterial hypertension. Eur Respir Rev 2021, 30 (162).

22. Neelakantan, S.; Xiang, Q.; Chavan, S.; Li, K.; Ling, X.; Dixon, R.; Sacks, M.; Vanderslices, P.; Avazmohammadi, R., Abstract 14303: Structural Remodeling in the Left Ventricular Myocardium Underlies Systolic Dysfunction in Myocardial Infarction. Circulation 2021, 144 (Suppl\_1), A14303–A14303.

23. Neelakantan, S.; Kumar, M.; Mendiola, E. A.; Phelan, H.; Serpooshan, V.; Sadayappan, S.; Avazmohammadi, R., Multiscale characterization of left ventricle active behavior in the mouse. Acta Biomaterialia 2023, 162, 240–253.

24. Mendiola, E. A.; Neelakantan, S.; Xiang, Q.; Merchant, S.; Li, K.; Hsu, E. W.; Dixon, R. A. F.; Vanderslice, P.; Avazmohammadi, R., Contractile Adaptation of the Left Ventricle Post-myocardial Infarction: Predictions by Rodent-Specific Computational Modeling. Annals of Biomedical Engineering 2022.

25. Avazmohammadi, R.; Hill, M. R.; Simon, M. A.; Zhang, W.; Sacks, M. S., A novel constitutive model for passive right ventricular myocardium: evidence for myofiber--collagen fiber mechanical coupling. Biomechanics and modeling in mechanobiology 2017, 16 (2), 561--581.

26. Babaei, H.; Mendiola, E. A.; Neelakantan, S.; Xiang, Q.; Vang, A.; Dixon, R. A. F.; Shah, D. J.; Vanderslice, P.; Choudhary, G.; Avazmohammadi, R., A machine learning model to estimate myocardial stiffness from EDPVR. Sci Rep-Uk 2022, 12 (1).

27. Mehdi, R. R.; Mendiola, E. A.; Sears, A.; Choudhary, G.; Ohayon, J.; Pettigrew, R.; Avazmohammadi, R., Comparison of three machine learning methods to estimate myocardial stiffness. In Reduced Order Models for the Biomechanics of Living Organs, 2023; pp 363--382.

28. He, Y.; Northrup, H.; Le, H.; Cheung, A. K.; Berceli, S. A.; Shiu, Y. T., Medical Image-Based Computational Fluid Dynamics and Fluid-Structure Interaction Analysis in Vascular Diseases. Front Bioeng Biotech 2022, 10.

29. Neelakantan, S.; Manning, E. P.; Zhang, P.; Choudhary, G.; Avazmohammadi, R., Right Ventricular Myocardial Stiffening is Associated With Pulmonary Arterial Stiffening in Pulmonary Hypertension. Circulation 2023, 148.

30. Avazmohammadi, R.; Mendiola, E. A.; Soares, J. S.; Li, D. S.; Chen, Z.; Merchant, S.; Hsu, E. W.; Vanderslice, P.; Dixon, R. A. F.; Sacks, M. S., A Computational Cardiac Model for the Adaptation to Pulmonary Arterial Hypertension in the Rat. Annals of Biomedical Engineering 2018, 47 (1), 138–153.

31. Avazmohammadi, R.; Hill, M.; Simon, M.; Sacks, M., Transmural remodeling of right ventricular myocardium in response to pulmonary arterial hypertension. APL Bioengineering 2017, 1 (1), 016105.

32. Rain, S.; Handoko, M. L.; Trip, P.; Gan, C. T. J.; Westerhof, N.; Stienen, G. J.; Paulus, W. J.; Ottenheijm, C. A. C.; Marcus, J. T.; Dorfmüller, P.; Guignabert, C.; Humbert, M.; MacDonald, P.; dos Remedios, C.; Postmus, P. E.; Saripalli, C.; Hidalgo, C. G.; Granzier, H. L.; Vonk-Noordegraaf, A.; van der Velden, J.; de Man, F. S., Right Ventricular Diastolic Impairment in Patients With Pulmonary Arterial Hypertension. Circulation 2013, 128 (18), 2016–2025.

33. Rain, S.; Andersen, S.; Najafi, A.; Schultz, J. G.; Bós, D. D. G.; Handoko, M. L.; Bogaard, H. J.; Vonk-Noordegraaf, A.; Andersen, A.; van der Velden, J.; Ottenheijm, C. A. C.; de Man, F. S., Right Ventricular Myocardial Stiffness in Experimental Pulmonary Arterial Hypertension Relative Contribution of Fibrosis and Myofibril Stiffness. Circ-Heart Fail 2016, 9 (7).

34. Jani, V.; Aslam, M. I.; Fenwick, A. J.; Ma, W. K.; Gong, H.; Milburn, G.; Nissen, D.; Salazar, I. M. C.; Hanselman, O.; Mukherjee, M.; Halushka, M. K.; Margulies, K. B.; Campbell, K. S.; Irving, T. C.; Kass, D. A.; Hsu, S., Right Ventricular Sarcomere Contractile Depression and the Role of Thick Filament Activation in Human Heart Failure With Pulmonary Hypertension. Circulation 2023, 147 (25), 1919–1932.

35. Mendiola, E. A.; Wang, E.; Leatherman, A.; Xiang, Q.; Neelakantan, S.; Vanderslice, P.; Avazmohammadi, R. In A Micro-anatomical Model of the Infarcted Left Ventricle Border Zone to Study the Influence of Collagen Undulation, Functional Imaging and Modeling of the Heart, Cham, 2023//; Bernard, O.; Clarysse, P.; Duchateau, N.; Ohayon, J.; Viallon, M., Eds. Springer Nature Switzerland: Cham, 2023; pp 34–43.

36. Zeinali-Davarani, S.; Wang, Y. J.; Chow, M. J.; Turcotte, R.; Zhang, Y. H., Contribution of Collagen Fiber Undulation to Regional Biomechanical Properties Along Porcine Thoracic Aorta. J Biomech Eng-T Asme 2015, 137 (5).

37. Lindeman, J. H. N.; Ashcroft, B. A.; Beenakker, J. W. M.; van Es, M.; Koekkoek, N. B. R.; Prins, F. A.; Tielemans, J. F.; Abdul-Hussien, H.; Bank, R. A.; Oosterkamp, T. H., Distinct defects in collagen microarchitecture underlie vessel-wall failure in advanced abdominal aneurysms and aneurysms in Marfan syndrome. P Natl Acad Sci USA 2010, 107 (2), 862–865.

38. Martin, C.; Pham, T.; Sun, W., Significant differences in the material properties between aged human and porcine aortic tissues. Eur J Cardio-Thorac 2011, 40 (1), 28–34.

39. Sahani, R.; Wallace, C. H.; Jones, B. K.; Blemker, S. S., Diaphragm muscle fibrosis involves changes in collagen organization with mechanical implications in Duchenne muscular dystrophy. J Appl Physiol 2022, 132 (3), 653–672.

40. Mehdi, R. R. R.; Neelakantan, S.; Wang, E.; Zhang, P.; Choudhary, G.; Avazmohammadi, R., Abstract P2008: Contractile Adaptation Of The Right Ventricular Myocardium In Pulmonary Hypertension. Circ Res 2023, 133 (Suppl_1), AP2008–AP2008.

41. Mair, K. M.; Johansen, A. K. Z.; Wright, A. F.; Wallace, E.; MacLean, M. R., Pulmonary arterial hypertension: basis of sex differences in incidence and treatment response. Brit J Pharmacol 2014, 171 (3), 567–579.

42. Ventetuolo, C. E.; Moutchia, J.; Baird, G. L.; Appleby, D. H.; McClelland, R. L.; Minhas, J.; Min, J.; Holmes, J. H.; Urbanowicz, R. J.; Al-Naamani, N.; Kawut, S. M., Baseline Sex Differences in Pulmonary Arterial Hypertension Randomized Clinical Trials. Ann Am Thorac Soc 2023, 20 (1), 58–66.

43. Walker, A. M.; Langleben, D.; Korelitz, J. J.; Rich, S.; Rubin, L. J.; Strom, B. L.; Gonin, R.; Keast, S.; Badesch, D.; Barst, R. J.; Bourge, R. C.; Channick, R.; Frost, A.; Gaine, S.; McGoon, M.; McLaughlin, V.; Murali, S.; Oudiz, R. J.; Robbins, I. M.; Tapson, V.; Abenhaim, L.; Constantine, G., Temporal trends and drug exposures in pulmonary hypertension: An American experience. Am Heart J 2006, 152 (3), 521–526.

